# Previous legume identity influence wheat rhizosphere microbial communities and grain protein content

**DOI:** 10.64898/2025.12.17.694934

**Authors:** Emmy L’Espérance, Vincent Poirier, Stéphanie Lavergne, Étienne Yergeau

**Affiliations:** Centre Armand-Frappier Santé Biotechnologie, Institut national de la recherche scientifique, 531 boulevard des Prairies, Laval, QC, H7V 1B7, Canada; Institut de recherche en agriculture et agroalimentaire, Université du Québec en Abitibi-Témiscamingue, 79 rue Côté, Notre-Dame-du-Nord, QC, J0Z 3B0, Canada

**Author notes:** Correspondance to: Etienne Yergeau, Centre Armand-Frappier Santé Biotechnologie, Institut national de la recherche scientifique, 531 boulevard des Prairies, Laval, QC, H7V 1B7, Canada.

**Keywords:** Plant-Soil feedback, Microbial communities, Rhizosphere, Depolymerization, crop rotation, Grain quality

## Abstract

In crop rotation systems, plant-soil feedback (PSF) effects can modulate nutrient cycling in soil, like organic carbon (C) and nitrogen (N) cycling. Organic nitrogen is present in plant and microbial necromass, and it must be depolymerized by microbes to become available for crops. Since plant identity can modulate microbial diversity and community composition, we thought that previous crop identity and its residue management would also change the microbial functional capacity, and thereby impact soil N availability, and the quality and yield of the following crop. To test this, two legumes (*Vicia faba* L., i.e. faba bean, and *Pisum sativum* L., i.e., yellow pea) were grown in two fields (Cloutier, and Palmarolle) in Abitibi-Témiscamingue, Québec, Canada (n=3 for each field). At the end of the growing season, we harvested peas and faba beans in both fields. During the following growing season, spring wheat (*Triticum aestivum* L.) was sown and received, or not granulated chicken manure as an organic fertilizer. We determined the diversity and composition of the microbial communities and their enzymatic depolymerization capacity in the soil and the rhizosphere each growing season. During wheat growth, previous legumes shaped bacterial (p-value = 0.006) and fungal (p-value = 0.001) communities without modulating the enzymatic activity of wheat-associated rhizosphere microbes. However, faba bean as a previous crop increased soil ammonium and wheat grain protein content at harvest as compared to peas at Cloutier. Altogether, our results show that faba beans can enhance wheat N nutrition, without a concomitant increase in potential protein or cellulose depolymerization, suggesting more mineralization due to increases in fungal: bacterial ratio or in the availability of substrates. Understanding plant-soil feedback in crop rotation systems is crucial to improve our practices and sustainably meet crops’ nutritional needs.

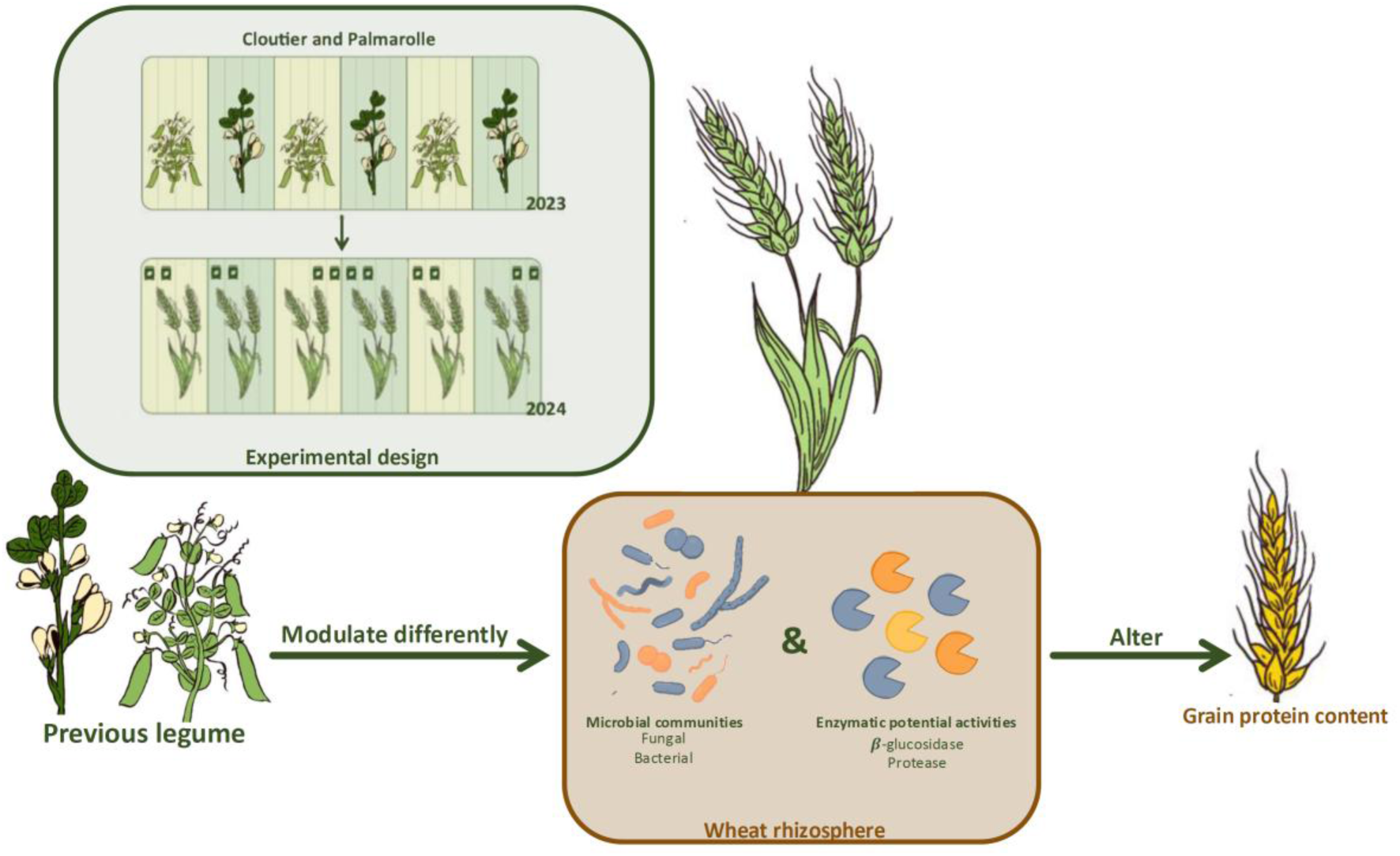

**Highlights:**

- Previous crop identity distinctively shaped wheat rhizosphere microbial communities
- Previous legume did not altered the enzymatic activity of wheat-associated rhizosphere microbes, but increased soil ammonium
- Wheat grain protein content was higher following faba bean than peas

## 1. Introduction

Agricultural practices have a significant impact on belowground microbial communities, altering their functionality, community structure, and diversity (Calderon et al., 2001; Li et al., 2020; Zhao et al., 2020). Monocropping often leads to lower soil microbial diversity and decreased plant growth and yield compared to crop rotation systems (Bak et al., 2022; Adebayo et al., 2025). Crop rotation systems (i.e., growing different crop species sequentially on the same land) enhance soil nutrient content and nutrient cycling (Wang et al., 2023c), limit the establishment of plant pathogens (Zhou et al., 2023) and enhance soil health and microbial diversity (Yang et al., 2024). Crop rotation systems are therefore fundamental in organic farming to maximize yield without the use of synthetic chemical inputs (Gamage et al., 2023).

The benefits of crop rotation systems can be explained by positive plant-soil feedback (PSF). The previous crop modulates soil microbial communities and physicochemical properties, impacting the next crop (Mariotte et al., 2018; Wang et al., 2025). Indeed, plants release a myriad of compounds, shaping spatially and temporally distinct microbial communities (Zhalnina et al., 2018). Positive plant-soil feedback includes pathogen reduction and beneficial microbes’ enrichment (Ma et al., 2017; Hannula et al., 2021). *Arabidopsis thaliana* changes its exudation pattern in response to the pathogenic bacterium *P. syringae* (Yuan et al., 2018). After six passages in the same soil, these changes created soil microbial communities antagonist to the pathogen, resulting in healthy plants (Yuan et al., 2018). Due to highly diversified plant traits, functions and exudation patterns, PSF effects are species-specific, meaning that we cannot associate a PSF effect with a plant family (Hannula et al., 2020). Plant-soil feedbacks are also not limited to the original plant species and can affect other plants grown in the same soil, creating multiple possibilities for crop rotation systems (Murrell et al., 2020). For instance, winter wheat (*Triticum aestivum* L.) grown continuously had a lower yield compared to winter wheat growing after oilseed rape, because oilseed rape changed the microbial communities and the abundances of functional genes related to inorganic N-cycling (Kaloterakis et al., 2025). This plant-specific rhizospheric effect can persist for many growing seasons (Bay et al., 2021).

Besides changing the microbial community structure and abundance of specific genes, crops also affect more generally the carbon (C) and nitrogen (N) cycles (Knops et al., 2002). Plant-soil feedback effects on nutrient cycling can be attributed to both living and dead crops. Plant litter origin (above vs belowground) and chemistry mediate the decomposition rate (Freschet et al., 2013), and microbial decomposer abundance and diversity over time (Bhatnagar et al., 2018). A classic use of PSF in agricultural settings is legume-cereal rotations. Legumes are well known to enhance soil N through biological fixation, and using them in crop rotation systems replenishes soil N without using chemical fertilizers (Gan et al., 2015). But not all legumes are identical. For instance, faba bean (*Vicia faba* L.) fixed 185 kg N ha^-1^ and forage peas (*Pisum sativum* L.) 165 kg ha^-1^, resulting in different amounts of C and N in their residues (Lupwayi and Soon, 2015). Living plants also influence microbial decomposer communities and their functionality through their C-rich root exudates. Since litter and root exudates composition differ among plant species, different crops in rotation systems will not have the same effect on soil nutrient cycling and on the next crop grown (Huo et al., 2017). Therefore, the choice of rotation systems design, especially the choice of crop species, is crucial to optimize PSF effects associated with nutrient cycling. The mechanistic link between shifts in microbial communities caused by the crop’s identity, soil nutrients availability, and the following crop’s growth and nutrition is, however, not clear, especially in organic agroecosystems. One possible explanation could be changes in organic N depolymerization.

Organic N depolymerization is a limiting step of the enzymatic cascade leading to mineralization and nitrification, producing inorganic N available for crops (Jan et al., 2009). The input of C-rich compounds, from plant litter and/or root exudates, steers up the C: N ratio, forcing microbes to mine for nitrogen (Jones et al., 2009; Kuzyakov and Xu, 2013) by depolymerizing soil organic matter (SOM) containing organic N, such as protein, chitin and peptidoglycan (Zhu et al., 2014). When their elemental demand is met, microbes will mineralize the excess N into inorganic form (ammonium and nitrate), usable by plants. Not to mention that small organic N forms (e.g. amino acids) are also usable by plants, highlighting the importance of N depolymerization for crop nutrition (Näsholm et al., 1998). Extracellular enzymes produced by microbial decomposers mediate soil N depolymerization. Microbes will release extracellular enzymes in soil, and their enzymatic activities will be induced by the presence of their substrate (Geisseler and Horwath, 2008). Nitrogen release from depolymerization depends on the microbial decomposer communities, soil physicochemical properties, and the organic substrate availability (Séneca et al., 2021), all of which are influenced by the identity of the plant and the crop rotation system. Increased N depolymerization may be one of the mechanisms responsible for the benefits of crop rotation systems. Yet, to our knowledge, few studies have tested that hypothesis under organic crop production conditions.

Here, we hypothesize that peas and faba beans, two legumes, have different lasting influences on the microbial community diversity and composition, changing the microbial depolymerization capacity, and thereby impacting soil inorganic N availability and quality and yield of wheat grown the following season. To test this hypothesis, we grew wheat with and without organic fertilizer in plots where peas or faba beans had been previously grown, under production conditions at two different fields in the Abitibi-Témiscamingue region (Québec Canada).

## 2. Materials and methods

### 2.1 Experimental design

We conducted a two-year field experiment in Abitibi-Témiscamingue (Québec Canada) on two farms under actual farm operating conditions, one in Palmarolle (48.665°N, 79.201°W) and in Cloutier (48.00297°N, 79.15°W). The goal of the experiment was to compare the effect of two legume crops (faba bean and peas) on the microbial communities and activities of wheat planted the following year. We also wanted to compare four fertilization strategies for wheat: granulated chicken manure (organic fertilizer), clover (*Trifolium repens*, cv. Huia) cover crop, fertilizer+cover crop, control without fertilization nor cover crop. We set up a split- plot experiment, with three blocks per field. In the first year (legume growing season), we defined randomly two plots (faba bean and peas) within each block. The following year (wheat growing season), we randomly defined two subplots (organic fertilizer and no fertilizer), and then two sub-subplots (clover cover crop and no cover crop). This resulted in 24 plots per field (48 total) (Fig. 1).

**Figure 1:**
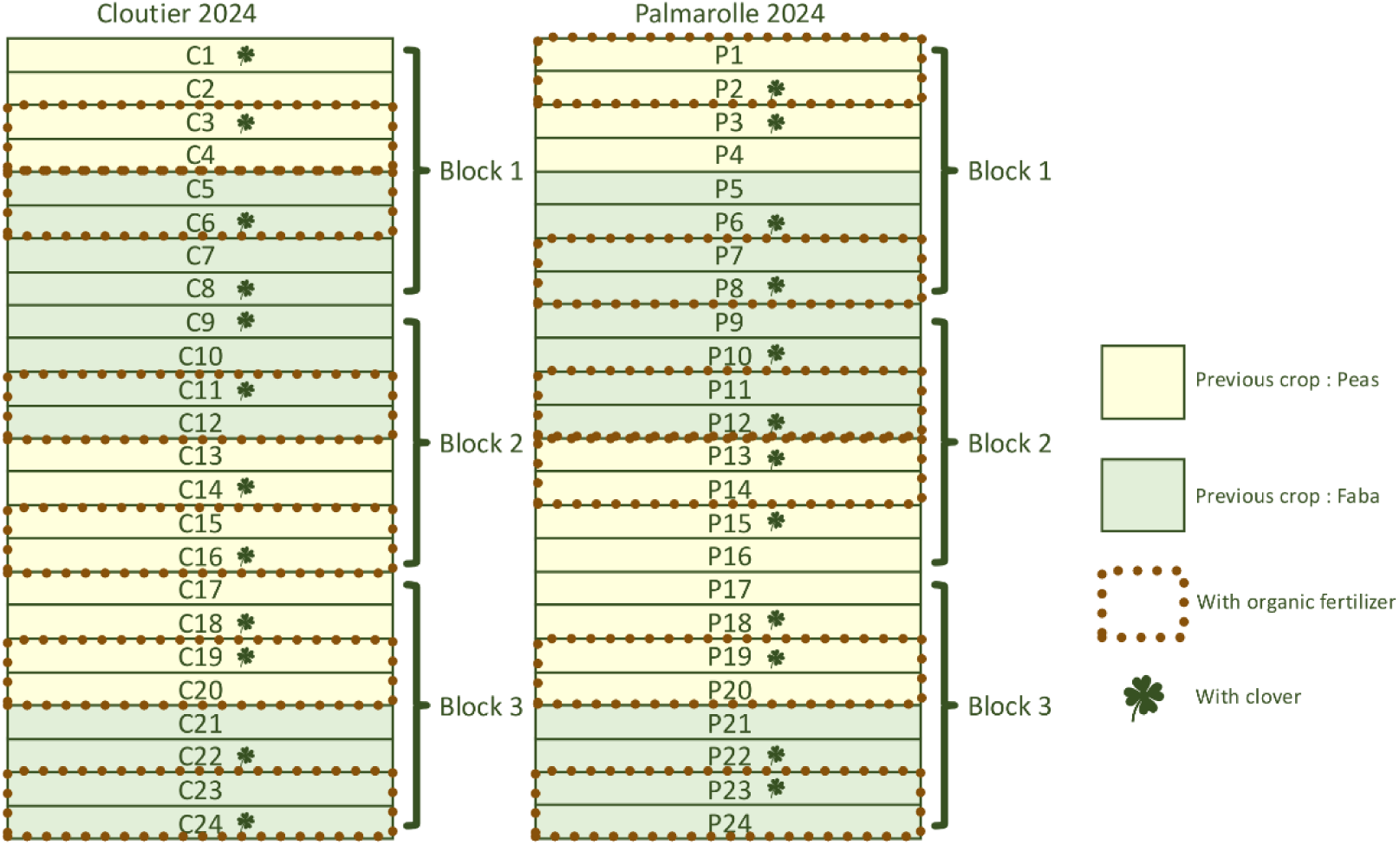
Experimental design of Cloutier and Palmarolle field in 2024. Colors (yellow and green) represent the previous crop sown in 2023. Brown dots line represents fertilized plots in 2024, and clovers represent plots that received clover as a cover crop. The cover crop was not used in our analysis, since it was not grown when we sampled mid-growing season (2024). Our experimental design allowed us to test the effects of previous crop, fertilization and wheat compartment on exoenzymes potential activities (Protease, deaminase and B-glucosidase), on inorganic N content and on the microbial community diversity and composition.

Both sites were grassland previously cropped to perennial forages for at least three years before our experiment. Palmarolle soil texture is clay with 80, 355 and 565 g kg^-1^ of sand (2000–50 µm), silt (50-20 µm) and clay (<2 µm) particles at 0-15 cm depth, respectively. It belongs to the Gleysolic order according to the World Reference Base for Soil Resource system (WRB, 2007) and the Canadian System of Soil Classification (Group et al., 1998). Cloutier soil texture is a silty clay with 38, 481 and 520 g kg^-1^ of sand, silt and clay particles at 0-15 cm depth, respectively, and also belongs to the Gleysolic order (Group et al., 1998) (WRB, 2007).

In fall 2022, both sites were moldboard plowed to a depth of 17 cm. In spring 2023, successive passes with a disk harrow and a field cultivator were carried out to prepare the seedbed. Faba beans and peas were sown in 2023, using the producer’s equipment (Table 1). Plots at Palmarolle were 11 m wide by 100 m long, while those at Cloutier were 4 m wide by 120 m long. At Palmarolle, faba bean was buried at the end of the growing season since it did not reach full maturity, while pea was harvested. At Cloutier, both crops were harvested at the end of the season (Table 1). But, since faba bean grains did not reach full maturity at Cloutier, residues were spread and buried a few days after harvest. Therefore, faba beans were used as green manure at both fields. Thus, residue management differed between both site (Cloutier: fragmented residues were buried a few days after and Palmarolle: residues were buried at the time of harvest).

**Table 1:** Sowing, Harvesting and sampling dates for Cloutier and Palmarolle in 2023 and 2024

| Field | Crops | Sowing rate (kg ha<sup>-1</sup>) | Sowing date | Harvesting date | Incorporation | Beginning of the season soil sampling date | Mid-growing season soil sampling date |
| --- | --- | --- | --- | --- | --- | --- | --- |
| Clou | Faba beans | 265 | 2023-06-07 | 2023-10-17<br>Buried afterward | Yes | 2023-06-06 | 2023-08-01 |
|  | Peas | 265 |  | 2023-09-27 | No |  |  |
|  | Wheat | 170 | 2024-06-04 | 2024-09-12 | No | 2024-05-23 | 2024-07-31 |
| Pal | Faba beans | 265 | 2023-05-21 | NA<br>Buried | Yes | 2023-05-21 | 2023-07-31 |
|  | Peas | 265 |  | 2023-09-18 | No |  |  |
|  | Wheat | 170 | 2024-05-30 | 2024-09-04 | No | 2024-05-21 | 2024-07-30 |

The following year (2024), the subplots were fertilized or not with the equivalent of 0.5 Mg N ha^-1^ and 1 Mg N ha^-1^ of granulated chicken manure (N:5-P:3-K:2) for Palmarolle and Cloutier, respectively. Wheat was sown on the same day as the fertilization occurred (i.e., May 30^th^ at Palmarolle and June 04 at Cloutier). The sub-subplots were assigned or not to receive clover seeds as a cover crop (sown on the same day and not incorporated into the soil). However, at the time of sampling, the clover had not established and we disregarded this treatment, treating the two sub-subplots from these treatments as replicates. At the end of the growing season, wheat was sampled and harvested (Table 1).

### 2.2 Soil sampling

Every year, we sampled twice during the season, in the beginning and mid-growing season (Table 1). Wheat was approximately in the tillering phase when we sampled in 2024. We sampled five points per sub-subplot (one in the center and four in the corners, at approximately 0.5 m from the sub-subplot borders), and we combined all samples into a composite, creating one composite sample for each of the 48 sub-subplots. From the five points, we sampled the bulk soil by collecting soil (0-15cm) between rows. When present (mid-growing season), we sampled the rhizosphere of the main crops (faba bean, peas or wheat) by gently removing plants from the same five points, shaking the roots to remove the excess soil and considering the remaining soil attached to the rhizosphere (i.e., the soil influenced by plant roots).

Soil samples were kept in a cooler until they were brought back to the laboratory. Samples were kept at 4 °C before sieving and aliquoting. We separated the rhizospheric soil from the roots before sieving. We sieved (4mm) rhizospheric and bulk soil, and we prepared two aliquots for each sample: one stored at 4 °C for enzymatic assays and ammonium/nitrate quantification, and another stored at -20 °C for DNA extraction.

### 2.3 Wheat sampling

At the end of the growing season, we sampled wheat to determine yield and protein content. Each sub-subplot was subsampled at four randomly distributed locations within the plot. At each location, we sampled a quadrat of wheat by placing two 76 cm long half-frames on the soil at about 45 cm width from each other, including 5 wheat rows (approximately 0.342 m^2^). At each subsample point, we measured the internal width between half-frames to calculate precisely the sampling area, and we harvested wheat. We stored wheat in paper bags and brought it back to the laboratory. Wheat was then dried at 35 °C for several days and kept for further analysis.

### 2.4 DNA extraction and sequencing

We extracted soil DNA from 0.25 g of bulk and rhizosphere soil (previously sieved at 4 mm) samples using the DNeasy PowerLyzer power soil kit (Qiagen) following the manufacturer’s protocol. DNA was eluted in 50 μL of elution buffer and was stored at −20°C until further analyses. We prepared libraries for 16S and ITS amplicon sequencing according to Illumina’s “16S Metagenomic Sequencing Library Preparation” guide (Part#15044223Rev.B). We prepared libraries for the bacterial 16 S rRNA gene using primers 16S: 515Y and 926R, and for the fungal ITS1 region, we used the primers ITS1F and 58A2R. NextSeq (Illumina) sequencing (PE300 10M reads) was performed at the *Centre d’expertise et de service Génome Québec* (CESGQ, Montréal, Canada).

### 2.5 Enzymatic potential activities quantification

We quantified the potential activity of three exoenzymes involved in SOM depolymerization (deaminases, β-glucosidases and proteases) using colorimetric methods. Every protocol follows the same principle: soil samples were incubated with an excess of substrate under optimal conditions. Therefore, we measured the maximum enzymatic capacity under optimal lab conditions, meaning that these results do not reflect real-time activity in the soil.

#### 2.5.1 Protease activity

To quantify the potential activity of soil protease, we used a protocol adapted from Jesmin et al. (2022). Briefly, we incubated 1 g of dried sieved (4 mm) soil with 2.5 mL sodium caseinate (2%) and 2.5 mL Tris buffer for two hours at 50°C and 100 rpm. After incubation, we added 5 mL Trichloroacetic acid (1 g L^-1^) to stop the enzymatic reaction, and we used 0.75 mL Na_2_CO_3_ and 0.25 mL Folin & Ciocalteu’s phenol (1:3) to quantify tyrosine, the product that produces a blue colour. We used 500 μL of every sample to measure the absorbance at 650-700 nm using a TECAN Infinite M plate reader. The potential protease activity is expressed in μmol tyrosine release g^−1^ dry soil h^−1^ incubation and was quantified using a tyrosine standard curve (2-fold serial dilution from 1.0 to 0.005 mM).

#### 2.5.2 β-glucosidases activity

To quantify the potential activity of soil β-glucosidase, we used a protocol adapted from Eivazi and Tabatabai (1988). Briefly, we incubated 1 g of dried sieved (4 mm) soil with a buffer solution containing p-nitrophenyl glycoside (25 mM) at 37°C for 1 h. After incubation, we added calcium chloride (0.5 M) and Tris buffer (0.1 M, pH 12) to extract p-nitrophenol, resulting from the degradation of the substrate. Extraction of the product produces a yellow colour, and we measured absorbance at 415 nm using a TECAN Infinite M plate reader. The potential activity is expressed in μmol p-nitrophenol released g^−1^ dry soil h^−1^ incubation and was quantified using a p-nitrophenol standard curve (2-fold serial dilution from 0.5 to 0.0078 g L^-1^).

#### 2.5.3 Deaminase activity

To quantify the potential activity of soil deaminases, we used a protocol adapted from Killham and Rashid (1986). Briefly, we incubated 1 g of dried sieved (4 mm) soil with 1.2-diamino-4-nitrobenzene (0.06 g L^-1^) for 20 h at 25 °C, 100 rpm. The substrate is red, and the colour intensity decreases with the deamination of the benzene ring. After incubation, we extracted twice the remaining substrate using methanol 100 % and filtered (Whatman no1) the solution to remove any impurities. We measured the absorbance at 405 nm using a TECAN Infinite M plate reader. The potential activity is expressed in and was quantified using a 1.2-diamino-4-itrobenzene standard curve (0/0.008/0.011/0.014/0.019/0.025/0.034/0.045/0.06 g L^-1^).

### 2.6 Available inorganic nitrogen quantification

To quantify ammonium and nitrate, we first needed to extract nitrogen from the soil, using a protocol adapted from Maynard et al. (1993).

To do so, we incubated 1 g of dried sieved (4 mm) soil with 10 mL of KCl (2 M) for 1 h at 25 °C at 100 rpm. After incubation, we filtered (Whatman no. 42) the soil slurry to remove any soil and debris. Soil extracts were kept at 4 °C until further analyses.

#### 2.6.1 Ammonium quantification

To quantify ammonium in soil extracts, we adapted a protocol from Rhine et al. (1998). Briefly, we added 100 μL of the sample (standard and/or soil extract) to a 96-well plate. Using a multi-channel pipette, we added 50 μL of citrate reagent, 50 μL of phenylphenol nitroprusside, 25 μL of buffered hypochlorite reagent and 50 μL of deionized water to each well and incubated 45 min at room temperature. Based on the Berthelot method, ammonium in soil extracts reacts with the buffered hypochlorite solution and the phenylphenol nitroprusside, producing a blue product. After incubation, we measured the absorbance at 660 nm using a TECAN Infinite M plate reader and the concentration of ammonium is expressed in mg g^-1^ of air-dried soil and quantified using a (NH_4_)_2_SO_4_ standard curve (2-fold serial dilution from 20 to 0.0195 mg L^-1^).

#### 2.6.2 Nitrate quantification

To quantify nitrate in soil extracts, we adapted a protocol from Hood-Nowotny et al. (2010). Briefly, we added 100 μL of the sample (standard and/or soil extract) to a 96-well plate. Using a multi-channel pipette, we added 100 μL of VCl_3_ solution and 100 μL of the Griess reagent mix (a solution containing 1:1 Griess solution 1 and Griess solution 2) to each well, and we incubated the plate for 1 h at 37 °C. Based on the VCL_3_/Griess method, nitrate in soil extracts is converted to nitrite with VCL_3_, and nitrite reacts with both Griess solutions to produce a pink color. After incubation, we measured absorbance at 540 nm using a TECAN Infinite M plate reader and the concentration of nitrate is expressed in mg g^-1^ of soil and quantified with a standard curve of KNO_3_ (2-fold serial dilution from 10 to 0.0195 mg L^-1^).

### 2.7 Wheat grain yield and protein content

At the end of the 2024 growing season, wheat was hand harvested (see section 2.3) to be threshed using a laboratory thresher (DL-350, Wintersteiger, Ried im Innkreis, Austria). Grains were then cleaned with a sample cleaner (MNL Pfeuffer, Kitzingen, Germany). Grain yield (Y_FH_, kg ha^-1^) was determined in both fields at the subsample level by weighing clean grain harvested for the given surface area sampled at each subsampling point (see section 2.3). Grain protein content (%) was determined only at Cloutier and was analyzed using a Near-infrared spectrometer (Infratec 1241, FOSS, Hillerød, Denmark). Wheat yield was reported on a 13.5 % relative humidity content (Y_13.5_, kg ha^-1^) as follows: Y_13.5_ = (Y_FH_ x (100 – W_FH_)) / (100 – 15). We used Y_13.5_ for our statistical analyses. We determined yield and protein content (when applicable) for each subsample harvested in the field (four subsamples), and we used the mean of the subsamples to perform our statistical analyses.

### 2.8 Bioinformatic analyses

We analyzed amplicon sequencing data (16S rRNA gene and ITS region) using the DADA2 pipeline in R to process raw reads. For 16S, we used truncLen (280,250) and maxEE (2,2) to filter and trim forward and reverse reads. We used the pooled method for removing chimeras, and we assigned taxonomy with the SILVA reference database. After the assignation, we removed chloroplasts and mitochondria from our dataset before further analysis. For ITS, we used truncLen (270,250) and maxEE (2,2) to filter and trim forward and reverse reads. We used the pooled method for removing chimeras, and we assigned taxonomy with the UNITE reference database.

### 2.9 Statistical analysis and data visualization

We performed Statistical analyses in R (version 4.5.1, *The R Foundation for Statistical Computing),* and we created figures using the package ggplot2. We divided and performed statistical analyses for each field (i.e., Cloutier and Palmarolle) and for each year (i.e., 2023 and 2024) separately. First, we performed Shapiro-Wilk tests to check the normality of the residuals and Bartlett tests to check the homogeneity of variances. When the assumptions were met, we performed a full factorial analysis of variance (ANOVA). For 2024, we performed ANOVAs for the effect of the previous legume crop (i.e., pea and faba bean), organic fertilization (i.e., with or without granulated chicken manure), soil compartment (i.e., rhizosphere and bulk soil), while including the split-split-plot design in the model. For 2023, we performed ANOVAs for the effect of the previous legume crop (i.e., pea and faba bean) and soil compartment (i.e., rhizosphere and bulk soil), while including the split-split-plot design in the model. We did this for potential enzymatic activities, soil ammonium and nitrate content. When the assumptions were not met, we first tried to transform our data (log or square root transformation), and if that did not work, we performed a Kruskal-Wallis non-parametric test, testing each factor independently. Specifically, in the 2024 second sampling, Cloutier nitrate did not meet the normality of residuals assumption, Cloutier ammonium and protease activity did not meet the homogeneity of variances assumption, and deaminase data did not meet both assumptions. Palmarolle deaminase data did not meet the normality of residuals assumption. In the 2024 first sampling, Cloutier nitrate did not meet the homogeneity of variances assumption. Palmarolle nitrate and ammonium did not meet both assumptions and deaminase data did not meet the homogeneity of variances assumption. Finally, in 2023 second sampling, Cloutier nitrate did not meet the homogeneity of variances assumption. Palmarolle nitrate did not meet both assumptions, Palmarolle ammonium did not meet the normality of residuals assumption and deaminase data did not meet the homogeneity of variances assumption. We performed Kruskal-Wallis nonparametric tests to assess the effects of previous crop, organic fertilization, the interaction between previous crop and fertilization and block on wheat yield (Y_13.5_, kg ha^-1^) and protein content. We performed Kruskal-Wallis nonparametric tests to assess the effects of previous crop, organic fertilization, soil compartment, and block on alpha-diversity and relative abundance data, since this data did not meet the assumption of parametric ANOVA. To analyze the bacterial and fungal community structure, we first transformed our dataset using the Hellinger transformation. We then used Euclidean distance matrices based on our transformed data to visualize the community structure using a principal coordinate analysis (PCoA). We first used betadisper to test the homogeneity of multivariate dispersion. We tested the effect of the previous crop, organic fertilization and soil compartment on the community using PERMANOVA (999 permutations). To consider the experimental design, we used the argument strata = block. To assess ASVs differently abundant in wheat plots after faba bean and peas, we performed three differential analysis (DEseq2, ANCOM-BC and LEfsE) and kept only ASVs found by all methods.

### 2.10 Data availability

The R code used to analyze the data and generate the figures is available on GitHub (https://github.com/le-labo-yergeau/). Raw sequencing reads were deposited in the NCBI SRA repository under BioProject accession PRJNA1390138.

## 3. Results

We focused our microbial analysis on the mid-growing season samples (2024) to assess previous legume crops’ PSF effects, except when it is mentioned.

### 3.1 Previous crop PSF on microbial communities’ composition

#### 3.1.1 Cloutier 2024

Before performing PERMANOVA, we tested the homogeneity of multivariate dispersion for both bacterial and fungal communities, and it was not significant, meaning that the results of the tests were not influenced by differential dispersion across treatments. The bacterial and fungal community compositions of wheat plots differed significantly depending on the previous legume sown in 2023 and the addition of organic fertilization (i.e. granulated chicken manure) in 2024 (Fig. 2.A and 2.C; Table S.1 and S.2). For the bacterial community (Fig. 2.A), there was also a significant interaction between the previous crop and the fertilization, meaning that depending on the previous crop sown, fertilization’s effect varied. In Figure 2.A, we can see that the addition of granulated chicken manure increases the variability of the bacterial community composition, especially when the previous crop was peas. Therefore, fertilization reduced the PSF effect of the previous crop. For the fungal community (Fig. 2.B), there was no interaction between the effect of the previous crop and the organic fertilization. The soil compartment (rhizosphere and bulk soil) modulated the fungal community composition (Fig. 2.B), but not the bacterial community.

**Figure 2:**
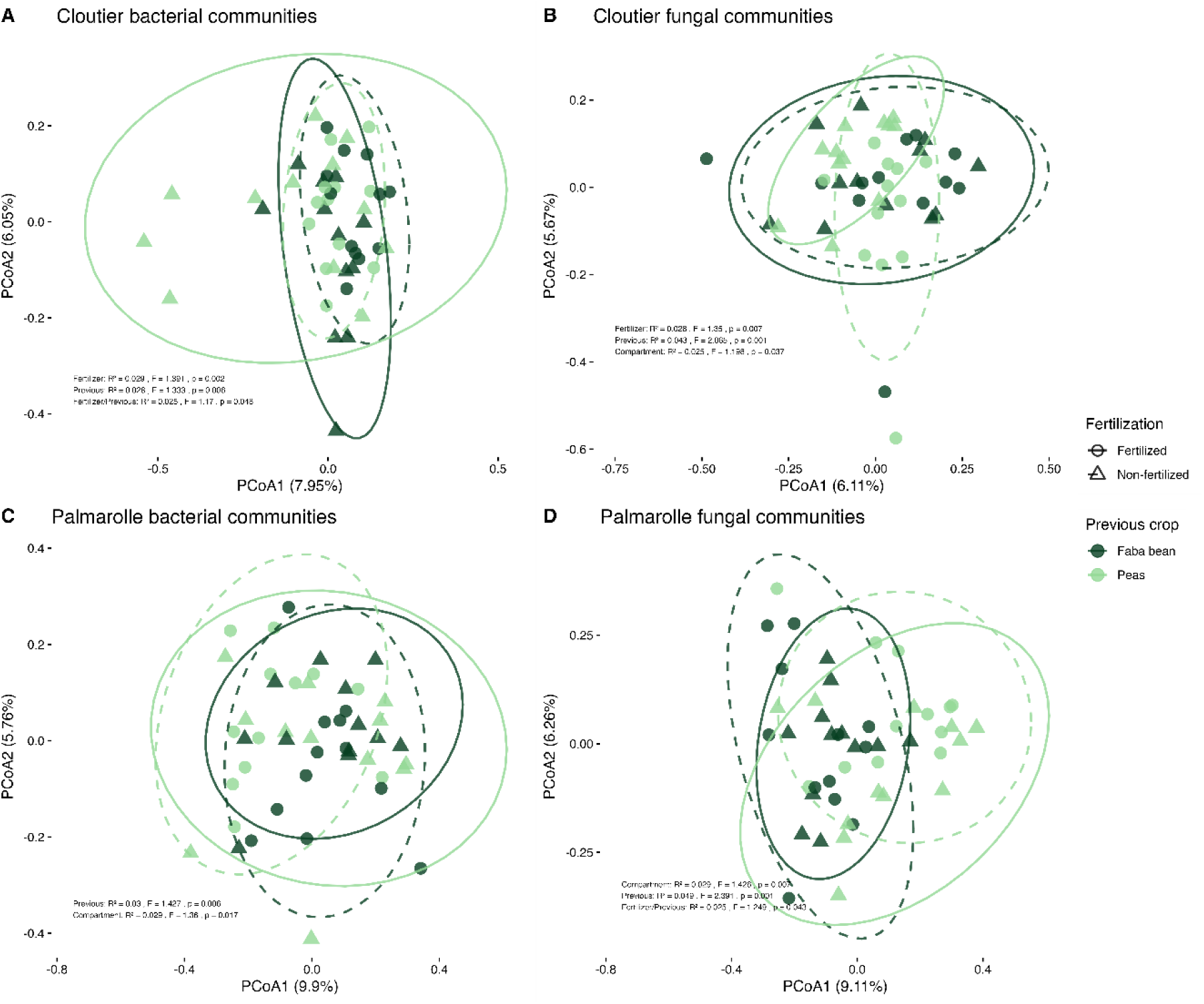
Beta diversity of the bacterial (16S) and fungal communities (ITS) from rhizosphere and bulk soil under wheat cropping at Cloutier (A, B) and Palmarolle (C, D) in 2024 using a Euclidean distance matrix and principal coordinate analysis. The effect of the previous crop (color) and organic fertilization (form and ellipse) is presented. For each field, all blocks are represented (n=3). The R^2^ and p-value correspond to the PERMANOVA results for these variables and the compartment (i.e., rhizospheric and bulk soil) variable.

After assessing that the previous legume crop identity shapes microbial communities in wheat plots differently, we tried to find which microbes’ relative abundance was different between plots that received faba beans and peas. After performing DESeq2, ANCOM-BC and LEfSe analysis, we did not find any bacterial ASV differentially abundant in wheat plots that received faba bean and peas (Fig. 3.B). On the other hand, we found 6 fungal ASVs differentially abundant in wheat plot depending on previous legume crops (Fig. 3.D; Table 2). Specifically, fungal ASVs # 16, 87, 387 and 686 were significantly more abundant in wheats plots that received peas as the previous legume crop and ASVs # 169 and 271 were significantly more abundant in wheat plots that received faba bean as the previous legume crop.

**Figure 3:**
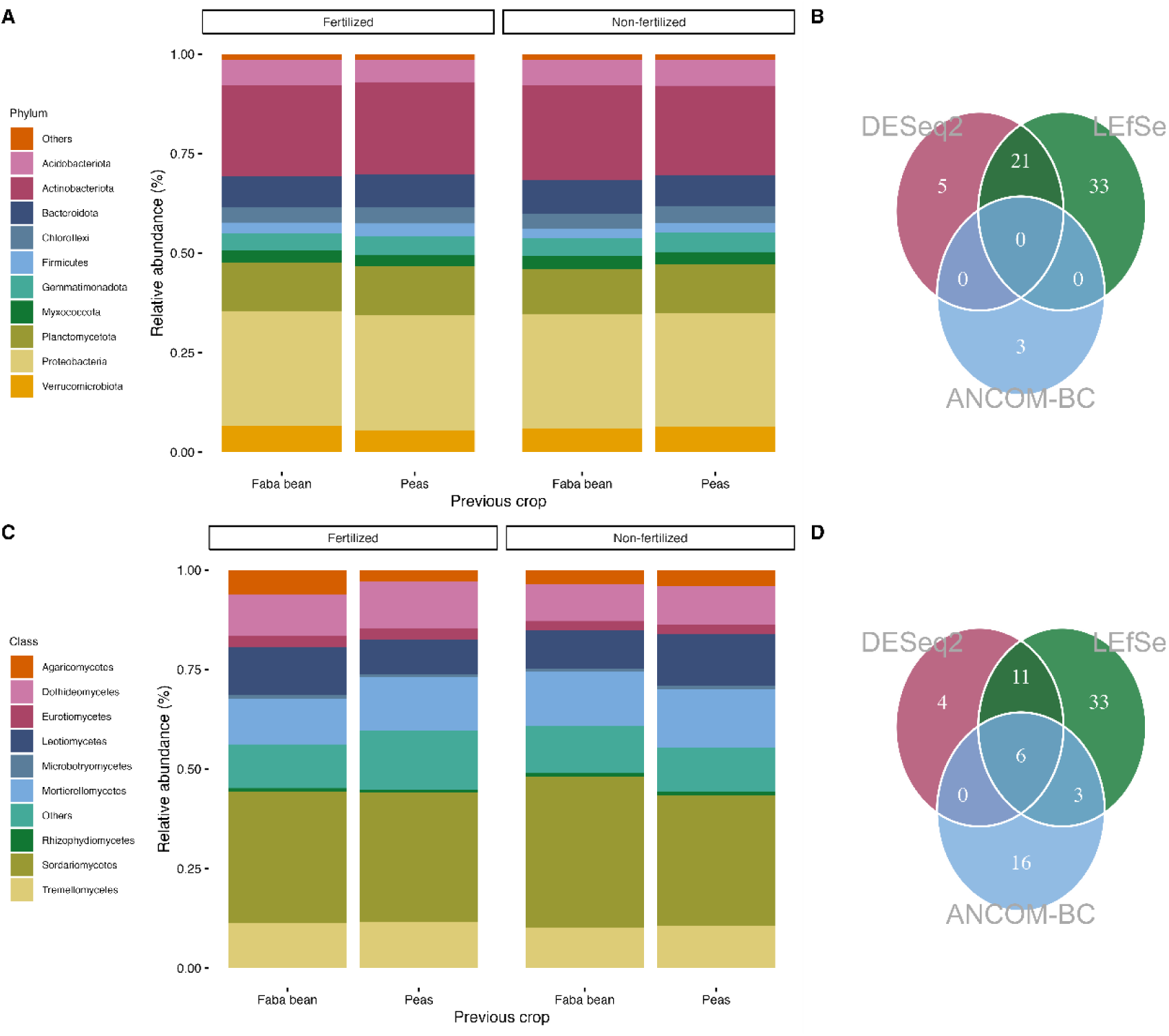
Compositions of the soil bacterial (A) and fungal (C) communities and Venn plot illustrating the number of differentially abundant bacterial (B) and fungal (D) ASV from rhizosphere and bulk soil under wheat cropping at Cloutier in 2024. Stacked bar charts contain the relative abundance of phyla for bacteria (A) and Classes for fungi. Venn plots illustrate the number of significantly differentially abundant ASVs found by DESeq2, ANCOM-BC and LEfSe analysis (p<0.05).

**Table 2:**
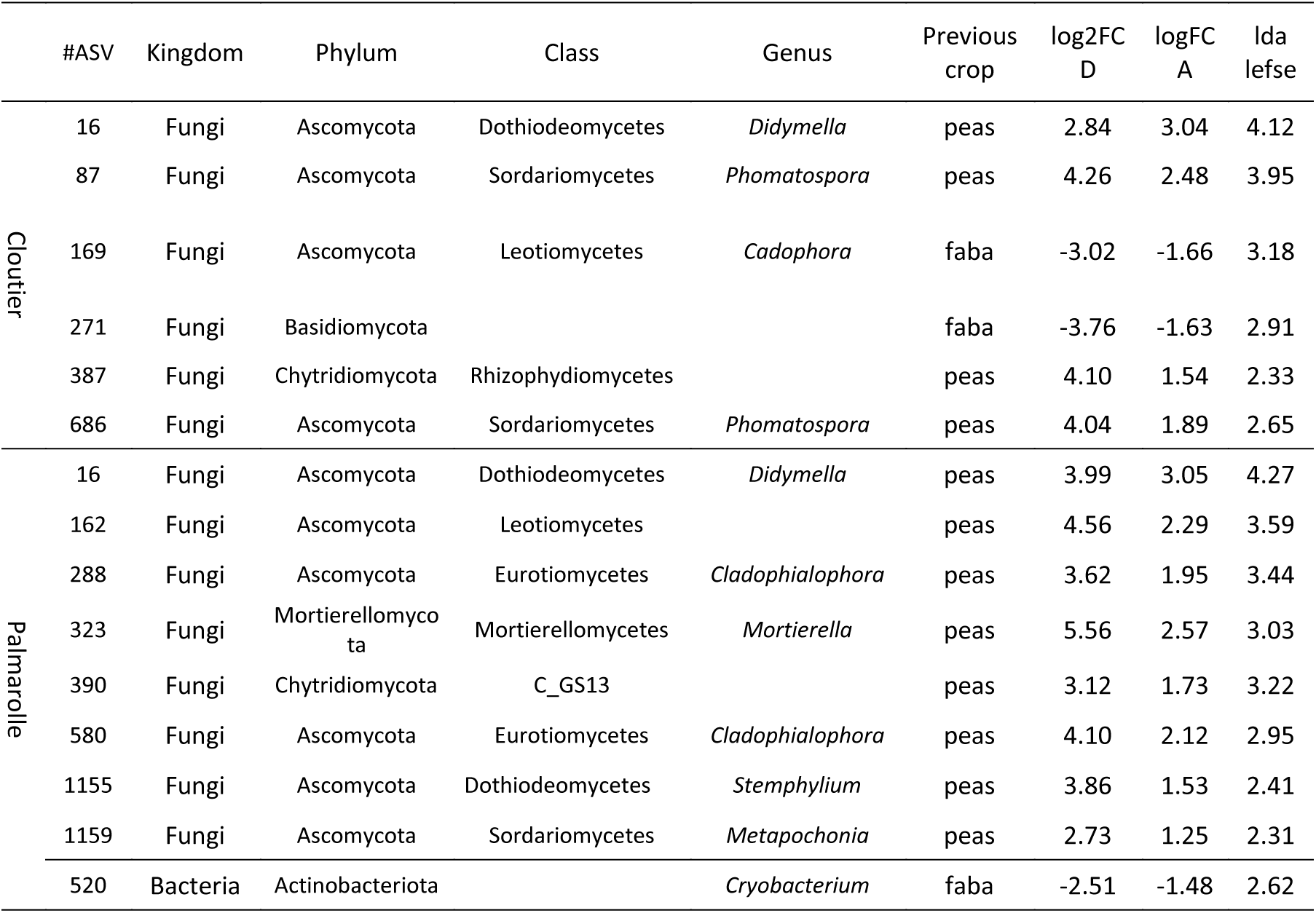
Bacterial and fungal ASVs differentially abundant in wheat plot at Cloutier and Palmarolle found with DESeq2, ANCOM-BC and LEfSe.

#### 3.1.2 Palmarolle 2024

Before performing PERMANOVA, we tested the homogeneity of multivariate dispersion, and it was not significant for fungi. However, we found significant differences for bacterial communities (p-value = 0.0249), which suggests that the PERMANOVA results could also be due to differential dispersion of the bacterial communities among treatments. The bacterial and fungal community composition of wheat plots is significantly different depending on the previous crop sown in 2023 and the soil compartment (Fig. 2.C and 2.D; Table S.1 and S.2). There is also a significant effect (p = 0.043) of the interaction between the previous crop and the fertilization on the fungal communities’ composition in Palmarolle (Fig. 2.D). Like the bacterial community composition in Cloutier, fertilization makes the fungal community composition more variable (Fig. 2.C).

After performing differential analysis (DESeq2, ANCOM-BC and LEfSe analysis), we found that the bacterial ASV # 520 was significantly more abundant in wheat plots that received faba bean as the previous legume crop (Fig. 4.B). We also found 8 fungal ASVs differentially abundant in wheat plots depending on previous legume crops (Fig. 4.D; Table 2). Specifically, fungal ASVs # 16, 162, 288, 323, 390, 580, 1155, 1159 were significantly more abundant in wheat plots that received peas as the previous legume crop.

**Figure 4:**
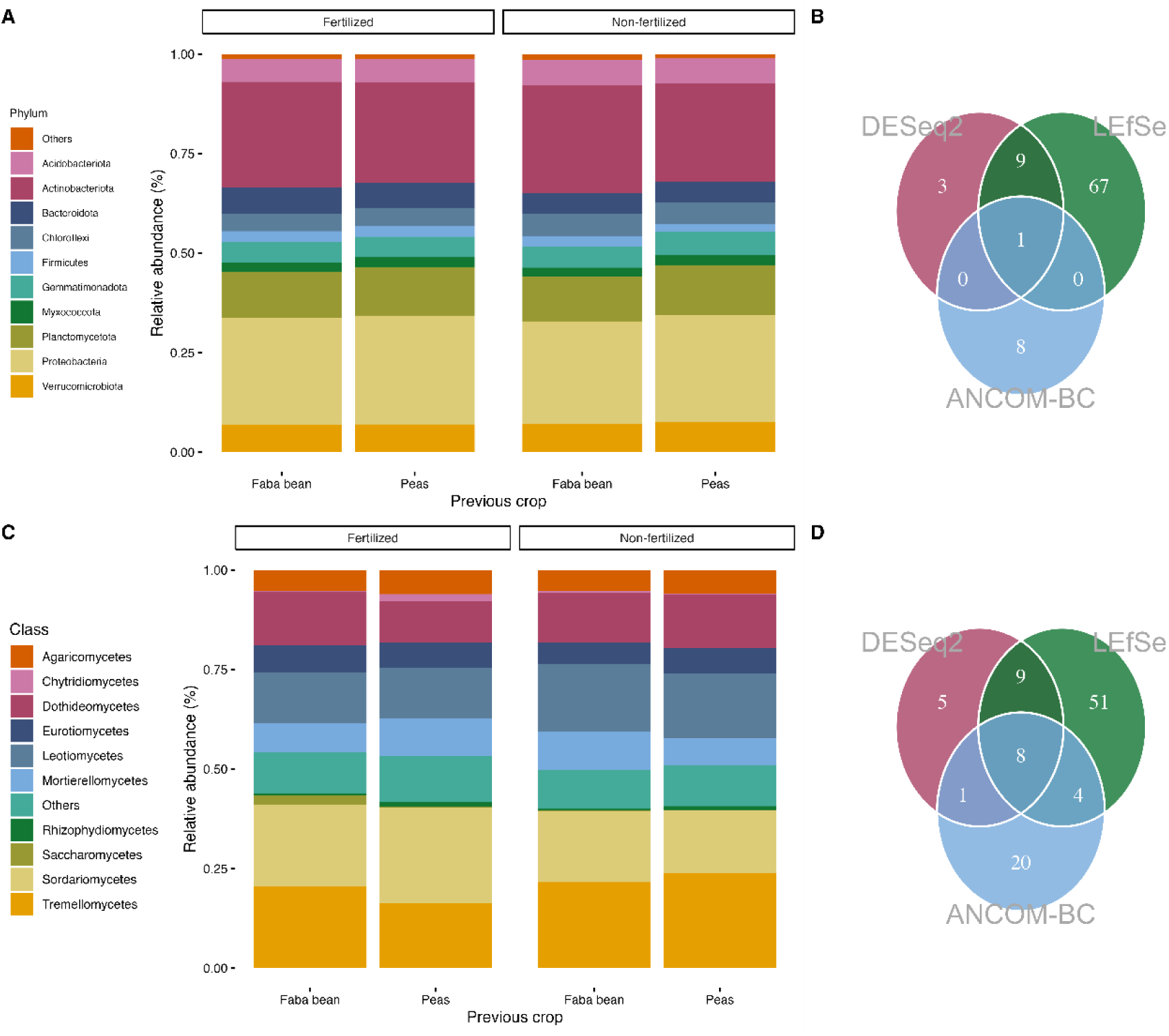
Compositions of the soil bacterial (A) and fungal (C) communities and Venn plot illustrating the number of differentially abundant bacterial (B) and fungal (D) ASV from rhizosphere and bulk soil under wheat cropping at Palmarolle in 2024. Stacked bar charts contain the relative abundance of phyla for bacteria (A) and Classes for fungi. Venn plots illustrate the number of significantly differentially abundant ASVs found by DESeq2, ANCOM-BC and LEfSe analysis (p<0.05).

### 3.2 Potential Enzymatic activities associated with depolymerization

#### 3.2.1 Cloutier 2024 and 2023

During wheat growth (2024 second sampling), there was significantly more β-glucosidase and deaminase potential activities in the wheat rhizosphere than in the bulk soil (Fig. 5.B and 5.C; Table 3). The potential β-glucosidase activity in wheat plots was higher, but not significant, after faba bean than after peas. The addition of organic fertilizer at the beginning of the season did not affect enzymatic activities during wheat growth. Before wheat growth (2024 first sampling), the potential protease activity was higher in wheat plots after peas than faba bean (Table 3). Fertilization did not affect enzymatic activities before wheat growth. In 2023, when faba bean and peas were growing, there was significantly more protease and deaminase potential activities in both legumes’ rhizosphere than in the bulk soil (Table 3).

**Figure 5:**
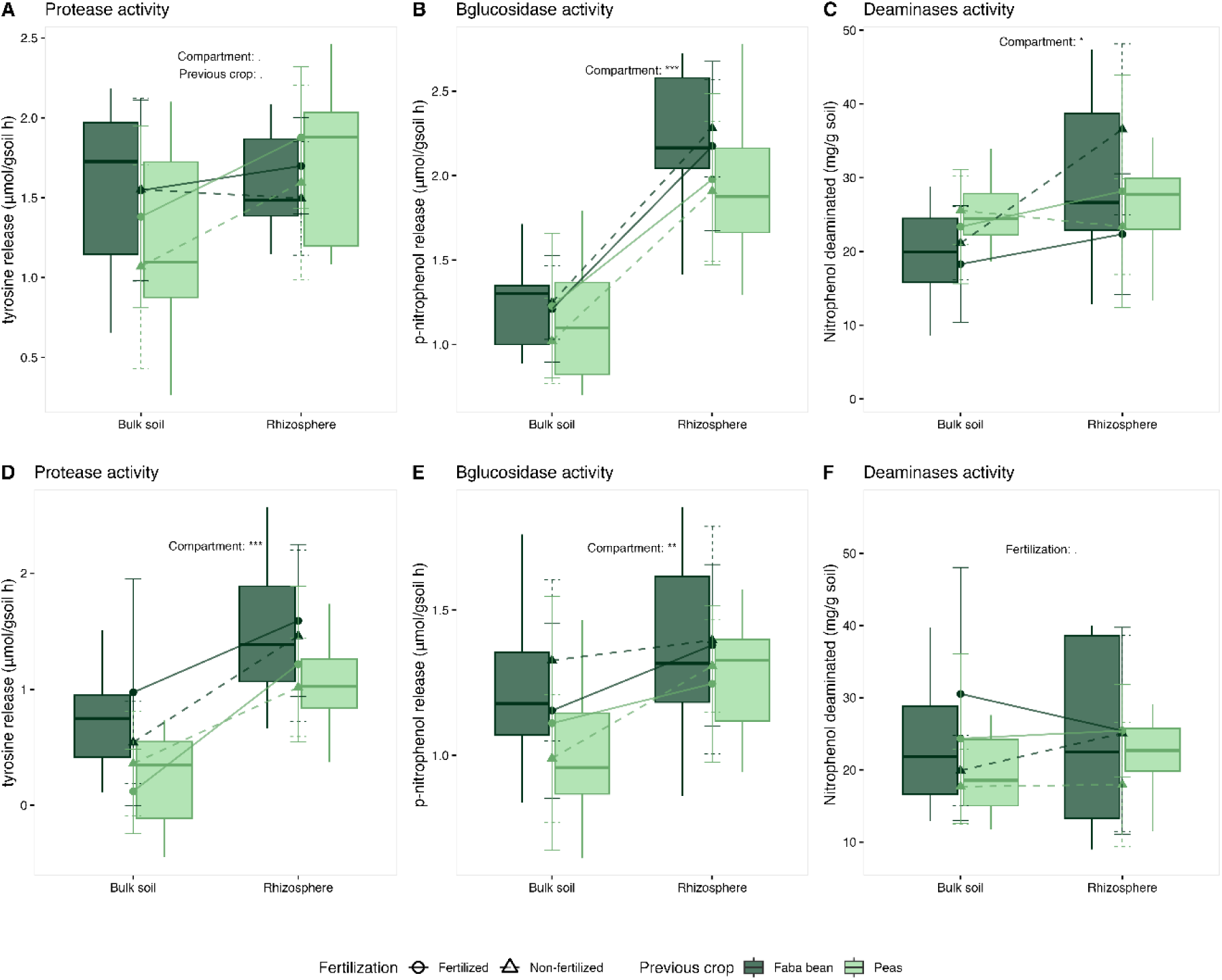
Protease, β-glucosidase and deaminases potential activities in the rhizosphere and bulk soil at 0-15 cm depth under wheat cropping at Cloutier (A, B, C) and Palmarolle (D, E, F) mid-growing season in 2024. The effects of the soil compartment, the previous crop, and fertilization are represented. The organic fertilization is represented by points (mean) and error bars (95% confidence interval) of both compartments and the previous crop. For each field and potential enzymatic activities, all blocks are represented (n=3). Significant results from Kruskal-Wallis or ANOVA tests for each enzyme’s potential activities (see Table 2) are identified with a *(p<0.05).

**Table 3:**
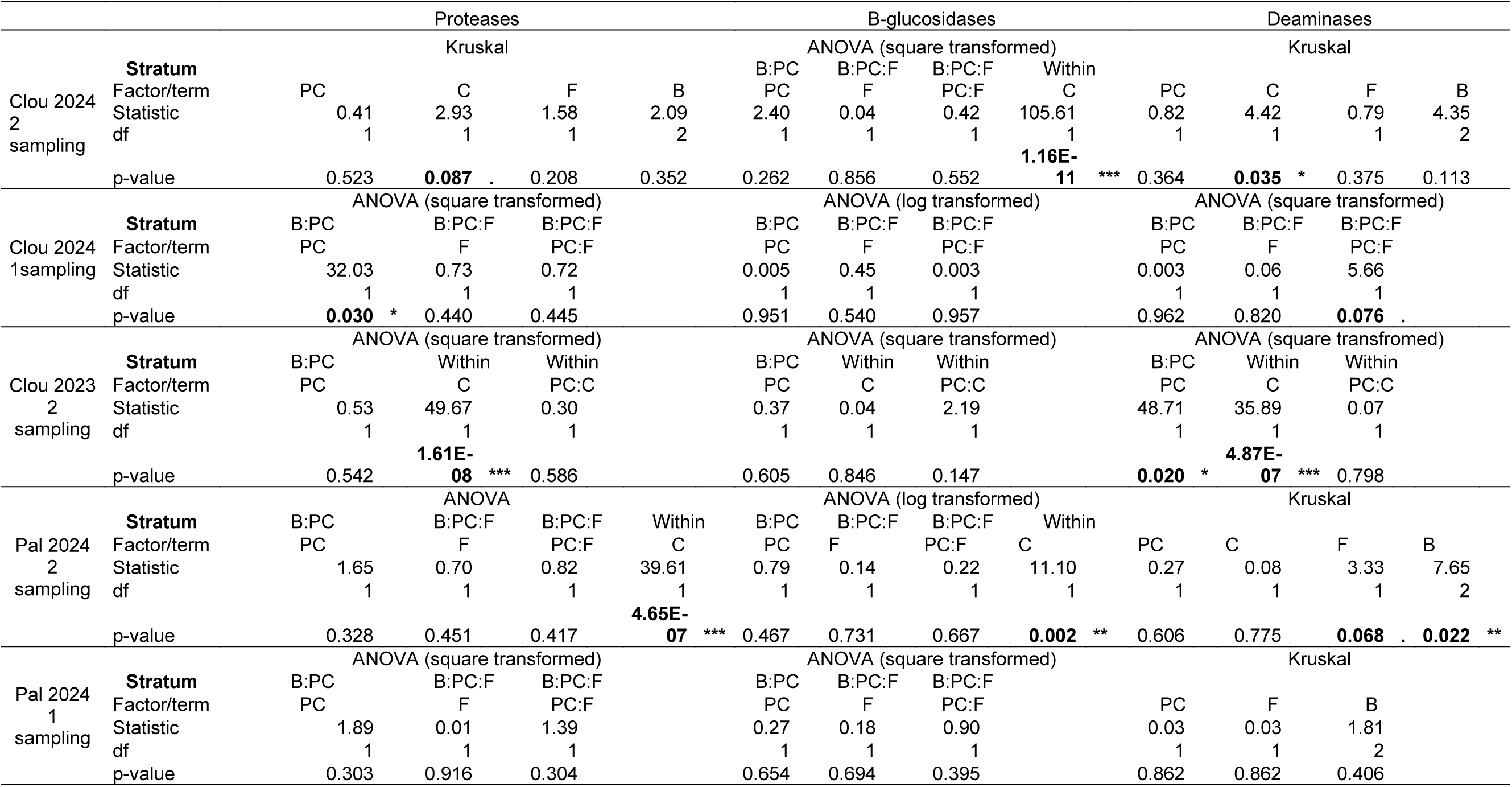
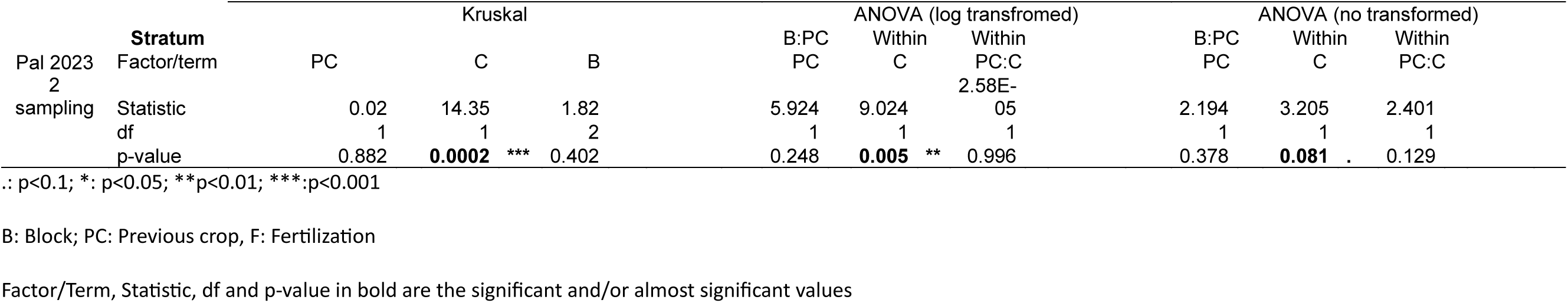
Effects of previous crop, organic fertilization and soil compartments (rhizospheric vs. bulk soil) on proteases, β -glucosidases and deaminases potential activities at 0-15 cm depth at Cloutier and Palmarolle during the wheat mid-season (2024 2 sampling), wheat beginning of the season (2024 1 sampling) and legumes mid-season (2023 2 sampling).

#### 3.2.2 Palmarolle 2024 and 2023

During wheat growth (2024 second sampling), there was significantly more protease and β-glucosidase potential activities in the wheat rhizosphere than in the bulk soil (Fig. 5.D and 5.E; Table 3). After faba bean, wheat plots had more protease and β-glucosidase activities than after peas. The addition of organic fertilizer at the beginning of the season did not affect enzymatic activities during wheat growth. Before wheat growth (2024 first sampling), fertilization and the previous crop did not influence the three exoenzymes potential activities tested. In 2023, when faba bean and peas were growing, there was significantly more protease, β-glucosidase, and deaminase potential activities in both legumes’ rhizosphere than in the bulk soil (Table 3).

### 3.3 Soil inorganic N content

#### 3.3.1 Cloutier 2024 and 2023

Figure 6 shows the amount of inorganic N in the bulk soil and the rhizosphere of wheat, at mid-growing season in 2024. In the wheat rhizospheric soil at Cloutier, there is significantly less inorganic N in ammonium and nitrate forms than in the bulk soil (Fig. 6.A and 6.B; Table 4). There is also a trend toward an increased concentration of ammonium (P=0.08) in the faba bean plots as compared to the pea plots (Fig. 6.A, Table 4). This could be related to the nearly significant (P= 0.084) compartment x previous legume interaction at Cloutier in 2023 (legume growing season) that stemmed from a higher concentration of ammonium in the rhizosphere vs. bulk soil, but only for faba bean (Table 4). In the first sampling of 2024, Cloutier nitrate concentration significantly differed between blocks. When legumes were the main crop (2023 second sampling), there was less nitrate in both legumes’ rhizosphere than in the bulk soil.

**Figure 6:**
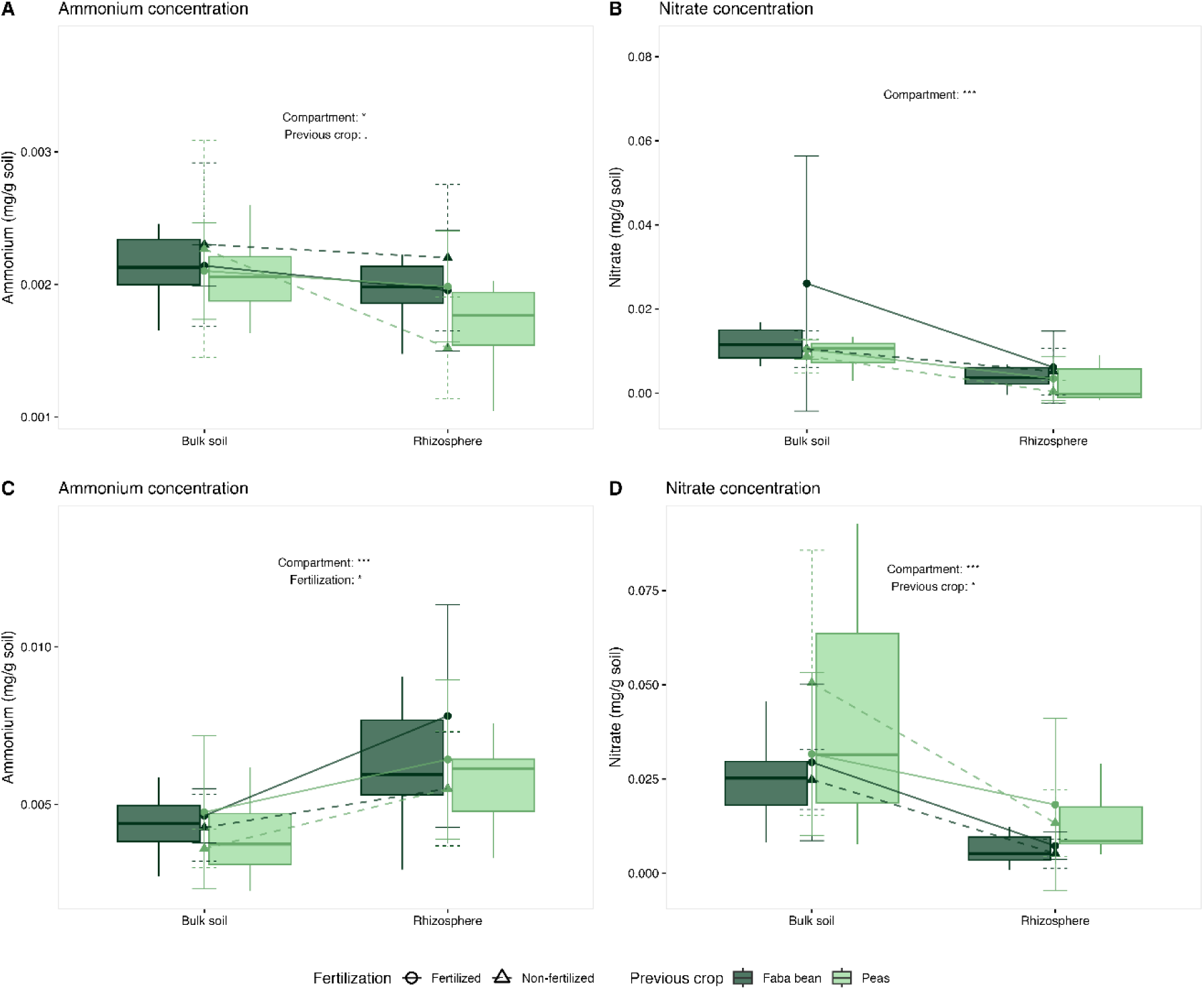
Wheat rhizosphere and bulk soil ammonium and nitrate content mid-growing season at Cloutier (A, B) and Palmarolle (C, D) at 0-15 cm depth. The effects of the compartment, the previous crop, and fertilization are represented. The organic fertilization is represented by points (mean) and error bars (95% confidence interval) of both compartments and the previous crop. For each field and N content, all blocks are represented (n=3). Significant results from Kruskal-Wallis or ANOVA tests for ammonium and nitrate content (see Table 3) are identified with a *(p<0.05).

**Table 4:**
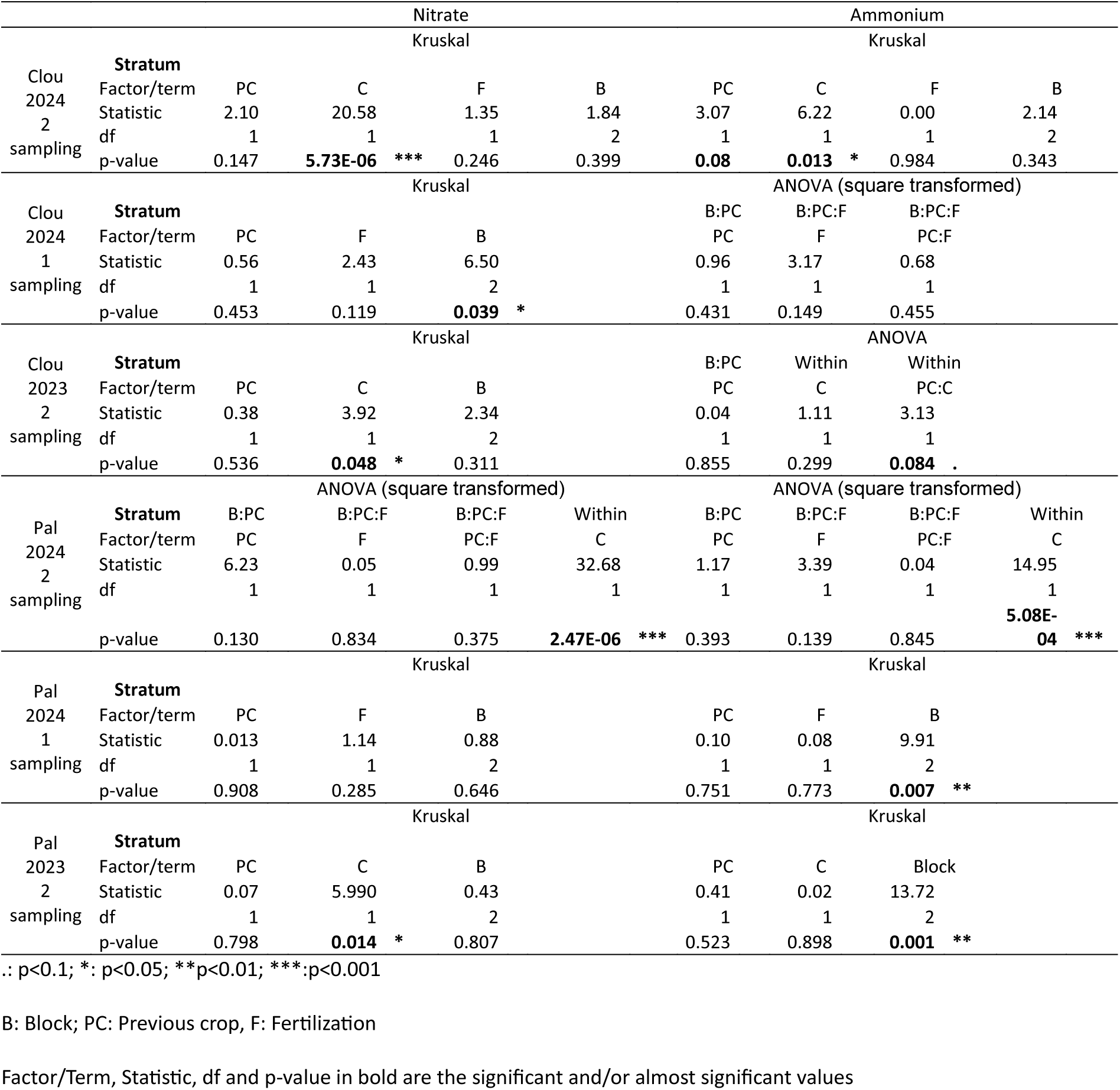
Effects of previous crop, organic fertilization and soil compartments (rhizospheric vs. bulk soil) on nitrate and ammonium content at 0-15 cm depth at Cloutier and Palmarolle during the wheat mid-season (2024 2 sampling), wheat beginning of the season (2024 1 sampling) and legumes mid-season (2023 2 sampling).

#### 3.3.2 Palmarolle 2024 and 2023

In the wheat rhizospheric soil at Palmarolle, there is significantly less nitrate, but more ammonium than in the bulk soil (Fig. 6.C and 6.D; Table 6). Also, at Palmarolle, there was less nitrate in wheat plots after faba beans than after peas, regardless of the compartment (Fig. 6.D). In the first sampling of 2024, Palmarolle ammonium concentration significantly differed between blocks. When legumes were the main crop (2023 second sampling), there was less nitrate in both legumes’ rhizosphere than in the bulk soil and ammonium concentration differed significantly between blocks.

### 3.4 Wheat yield and grain quality

#### 3.4.1 Cloutier 2024

The addition of organic fertilizer significantly enhanced wheat grain protein content (Fig. 7: Table 5). Also, faba bean, when used as the previous crop, significantly increased wheat grain protein content as compared to peas (Fig. 7).

**Figure 7:**
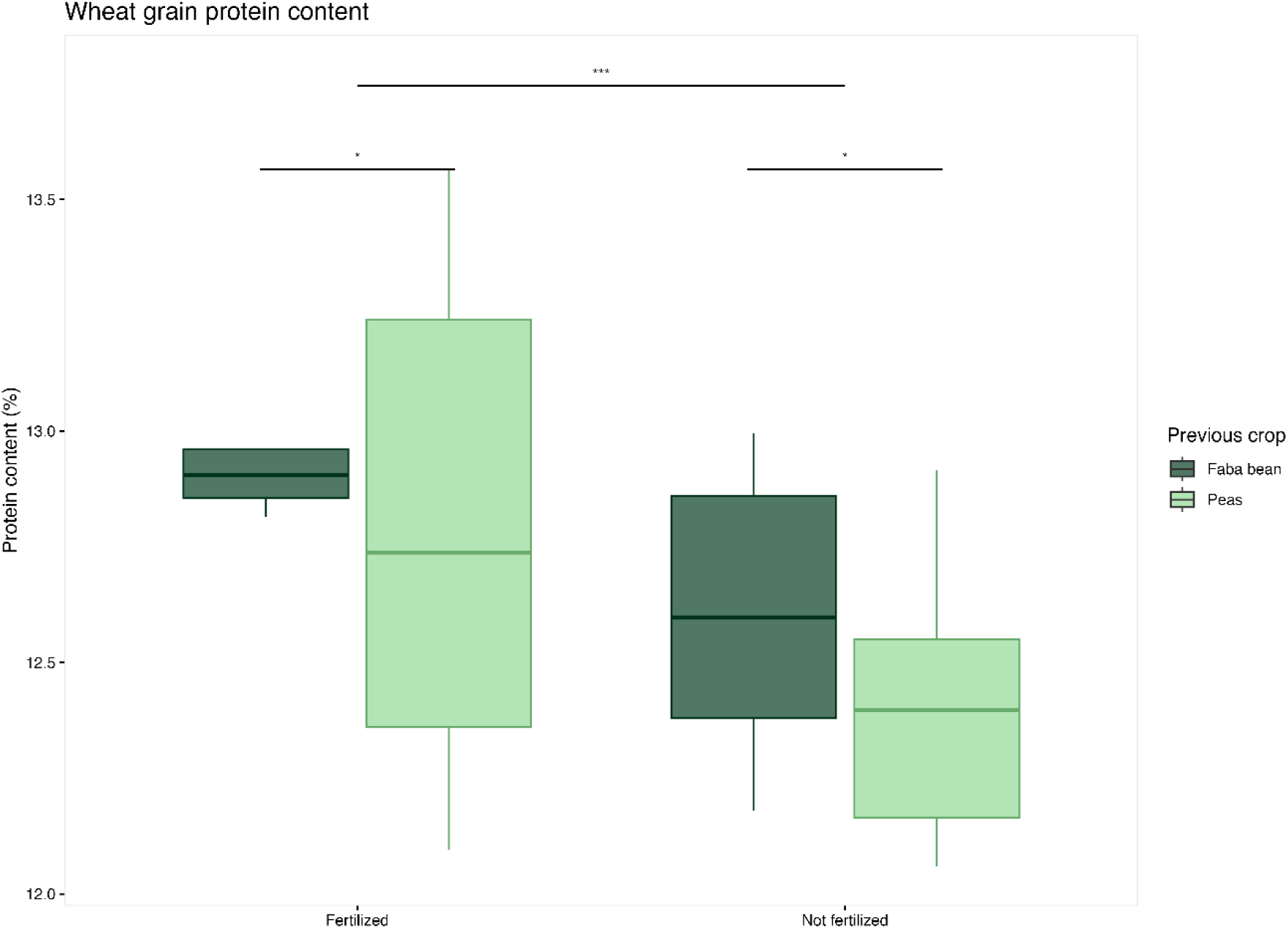
Wheat grain protein content of each organic fertilization treatment and previous crops after harvest at Cloutier. The effects of the previous crop and fertilization are represented. Significant results from Kruskal-Wallis tests (see Table 3) are identified with a **(p<0.05).

**Table 5:** Kruskal-Wallis results for the effect of organic fertilization and previous crops on grain protein content and yield (Y_13.5_) at Cloutier

| Response variable | Factor | Statistic | p-value | p.adj |  |
| --- | --- | --- | --- | --- | --- |
| Yield(kg ha<sup>-1</sup>) | Fertilization | 0.093 | 0.7609 | 0.7609 |  |
|  | Previous crop | 1.245 | 0.2645 | 0.3174 |  |
|  | Fertilization :<br>Previous crop | 7.265 | 0.0639 | 0.1024 |  |
|  | **Block** | **10.619** | **0.0050** | **0.0148** | **\*** |
| Protein content (%) | **Fertilization** | **14.087** | **0.0002** | **0.0007** | **\*\*\*** |
|  | **Previous crop** | **5.927** | **0.0149** | **0.0298** | **\*** |
|  | **Fertilization :**<br>**Previous crop** | **20.106** | **0.0002** | **0.0006** | **\*\*\*** |
|  | **Block** | **10.048** | **0.0066** | **0.0158** | **\*** |

\*: p<0.05; \*\*p<0.01; \*\*\*:p<0.001
p.adj refers to the p-value with Benjamini-Hochberg correction
Factor/Term, Statistic, df, p-value and p.adj in bold are the significant values

For Palmarolle, we were only able to determine the grain yield at the end of the growing season. We found no significant effect of fertilization and the previous crop on grain yield (Table 5).

**Table 5:** Kruskal-Wallis results for the effect of organic fertilization and previous crops on grain yield (Y_13.5_) at at Palmarolle

| Response variable | Factor | Statistic | p-value | p.adj |
| --- | --- | --- | --- | --- |
| Yield (kg ha<sup>-1</sup>) | Fertilization | 0.257 | 0.6120 | 0.9180 |
|  | Previous crop | 0.833 | 0.3613 | 0.7225 |
|  | Fertilization :<br>Previous crop | 1.461 | 0.6913 | 0.9217 |
|  | Block | 0.247 | 0.8838 | 1.0000 |

\*: $p < 0.05$ ; \*\*: $p < 0.01$ ; \*\*\*: $p < 0.001$
p.adj refers to the p-value with Benjamini-Hochberg correction
Factor/Term, Statistic, df, p-value and p.adj in bold are the significant values

## 4. Discussion

This study aimed to evaluate the impact of the identity of the previous legume crop on microbial community composition and functional capacity in the following growing season. We hypothesized that a part of the positive legume-cereal rotation effect on wheat yield and grain quality could be mediated by changes in the C and N depolymerization capacity of the microbial community. First, we confirmed that legume identity modifies soil microbial communities in the following growing season, and that PSF effects can be transferred to other plant species (in our case, from legumes to wheat). We also found, at Cloutier, that faba bean as previous crop increased soil ammonium content and wheat grain protein content, a key parameter of grain baking quality (Nuttall et al., 2017). We did not, however, find significant changes in potential protein and cellulose depolymerization activities, infirming our main hypothesis. Instead, the mechanism behind the effect of PSF on wheat grain quality could be an increased mineralization linked to shifts in the relative dominance of bacteria and fungi or a higher substrate availability in faba bean plots.

Under real farming conditions in fields located 100 km apart, we confirmed that the previous crops’ PSF influenced the composition of microbial communities. In fact, among all factors tested, the previous crop was the only one that consistently shaped microbial communities in both fields. This highlights the strength of the previous crop effect on microbial communities. Indeed, due to logistics (e.g., the size of plots needed to match the seeder and the fertilizer applicator) and field size, we could not set up more than three complete blocks. Additionally, there was quite some variation between the blocks, but we still found significant evidence of a PSF effect. Previous crop PSF effects on microbial community composition has been observed before (Wang et al., 2023a; Dang et al., 2024). We also found ASVs significantly differently abundant depending on the previous crop identity, proving the legacy effect of previous legume crops on microbial communities. In both fields, fungal communities were more altered by the previous crop identity and residue management than bacterial communities. Fungi are known to be key players in decomposition and can form close relationships with plants, as symbionts and pathogens (Zeilinger et al., 2016). Therefore, changes in fungal community composition and relative abundance due to the previous crop identity are expected (Li et al., 2023). Fungi are also known to be key players in plant compounds degradation, especially lignin and cellulose (Schneider et al., 2012; Arcand et al., 2016). For instance, in Cloutier’s wheat plots after faba bean, Leotiomycetes, more specifically *Cadophora luteo-olivacea,* were more abundant. *Cadophora spp.* possess a wide variety of exoenzymes associated with C and N depolymerization, such as protease and carbohydrate active enzymes (Bruyant et al., 2024). They could have potentially played a role in faba bean residue depolymerization and wheat nutrition. In Palmarolle’s wheat plots after faba bean, *Cryobacterium spp.* was more abundant. *Cryobacterium spp.* is often correlated to higher lignin and C content (Shamshitov et al., 2025). The incorporation of faba bean residue could have enhanced *Cryobacterium spp.* abundance during the wheat growing season. Fungi have a higher C:N ratio than bacteria and increases in fungal dominance in depolymerization processes would result in higher mineralization rate of the depolymerized organic N (Strickland and Rousk, 2010), without necessarily changing potential enzymatic activities.

Faba beans and peas are two legumes often used in crop rotation systems to replenish soil N pools. Both crops fix atmospheric N through a symbiosis with rhizobia (Jensen et al., 2010). Their PSF effects on microbial community composition were, however, distinct, supporting other findings (Yixuan et al., 2025). Faba bean is particularly efficient in increasing soil N for the following crop (Herridge et al., 2008; Peoples et al., 2009). Faba beans have a coarser root system (i.e., higher mean root diameter and lower specific root length) and allocate a higher proportion of total plant biomass belowground than pea (Guerrero-Ramírez et al., 2021), therefore adding more organic matter to the soil (Nuruzzaman et al., 2006). Differences in root system sizes can also alter soil physical properties, therefore changing microbial communities. With a root system four times bigger than peas, faba beans promote aggregate formation, creating different microenvironments for microbes in the next growing season (Oliveira et al., 2019). At Cloutier, we also found higher ammonium concentration in the rhizosphere of faba beans as compared to adjacent bulk soil, while there was no difference for peas. This difference between faba beans and peas was maintained during the wheat growing season, which could explain the higher grain protein content of wheat that followed faba beans. Indeed, even with similar potential depolymerization activities, the soil with more substrate (i.e. faba bean plots) will mineralize more N, as soil depolymerization activities are substrate-limited (Greenfield et al., 2020).

Fertilization alters soil properties and microbial communities (Luo et al., 2018), and it was expected that organic fertilization would affect microbial communities in our study. Interestingly, microbial communities from wheat growing season did not respond the same way to organic fertilization in both fields. It is important to note that the organic fertilizer amount was not the same in both fields, which could in part explain the difference between fields (Zhou et al., 2021). Nevertheless, organic fertilization increased the variability of microbial communities coming from peas as compared to faba beans. It suggests that faba beans have a stronger PSF effects due to a larger addition of root-derived organic matter and, incorporation of aboveground biomass. It creates a resource-rich environment, priming the microbial community to mineralize organic fertilizer and making it less susceptible to changes caused by fertilizer addition (Aschi et al., 2017).

The only significant difference in potential enzymatic activity we found between the two legumes was for protease activity that was higher in plots that received peas at Cloutier early in the season, before wheat was sown. In contrast to our main hypothesis, this increased protease activity did not seem to be beneficial for the wheat, as the wheat grains harvested from the pea plots had in fact a lower protein content than the ones harvested in the faba bean plots. This increase in protease potential activity, early in the season and in the absence of the crops, probably led to microbial immobilization of the depolymerized N. In that case, the immobilized N would have needed to be mineralized or re-mobilized from the microbial biomass to reach the wheat. Other enzymes that depolymerize chitin or peptidoglycan, two abundant soil organic N sources that make fungal and bacterial cell walls, respectively, could have been good targets to test this idea. However, since we observed less protein content in wheat grains from the pea plots, the N produced from protease activity probably remained immobilized. This suggest that the timing of depolymerization activities is paramount for it to be beneficial to crops. In wheat, the N assimilated by the plant is mostly taken up before and during the anthesis stage (Cox et al., 1985; Sharma et al., 2023). Our mid-season sampling might have missed the potential increase in depolymerization activities that could have increased wheat N uptake at key growth stages, resulting in higher protein content at the end of the season at Cloutier. Alternatively, as mentioned above, enzymatic activities are substrate-limited, and changes in potential enzymatic activities measured in the laboratory with excess substrate might not represent activities in the field.

Almost every exoenzyme’s potential activities were higher in the wheat rhizosphere than in the bulk soil, suggesting that wheat, through its root exudates, selects for a community proficient in depolymerization. Our results align with the rhizosphere priming theory, a plant competitive strategy for N (Kuzyakov and Xu, 2013; L’Espérance et al., 2024). Plants change the rhizosphere C: N ratio through their C-rich exudates to force microbes to depolymerize soil organic matter to scavenge for N. When the C-rich exudates are all consumed, microbes will mineralize the N excess, providing inorganic N to plants (Zhu et al., 2014). To confirm this hypothetical rhizosphere priming effect, future studies should measure rhizosphere C: N ratios and root exudates.

Unlike previous studies (Li et al., 2015; Lupwayi et al., 2019), the addition of organic fertilizer did not have a significant effect on the enzymatic activities, except for the deaminase activity at Palmarolle. This is surprising, since we added a substrate that can stimulate microbes to produce and release exoenzymes. In contrast, Breza et al., (2023) found that the addition of granular urea in more complex rotation systems disturbs the organic N cycle, reducing the N depolymerization rate to a similar rate as monocropping. In both studies, fertilization affects exoenzymes activities mediating N depolymerization (Breza et al., 2023). In our case, the lack of fertilization effects can be due to the fertilizer type, the application rate and/ or the sampling date. We chose to use a fertilizer used in this region for organic farms, and the equivalent of only 25-50 kg N ha^-1^ was applied. The amount applied – which is less than the recommended amount for spring wheat under organic management (∼100 kg N ha^-1^) – and type of fertilizer was chosen to accommodate farmers who must reconcile crop needs and fertilizer prices. The lack of fertilization effects can also be due to the sampling date. Our mid-growing season sampling point in 2024 was approximately two months after the application of the organic fertilizer. Organic fertilizer availability depends on the soil mineralization rate and usually takes longer to be available for plants and microbes (Wang et al., 2023b). However, Lazicki et al. (2020) showed that, under laboratory conditions, poultry manure-based granulated fertilizer was already mineralized after a few weeks of incubation. Therefore, nutrients derived from organic fertilizer were probably already immobilized in microbial or wheat biomass at the time of sampling, which would explain the lack of a fertilizer effect on enzymatic activities and soil ammonium and nitrate content.

Conducting a two-year field experiment under actual farm operating conditions is a daunting task, and differences between fields are unavoidable. Pedoclimatic conditions, residue management, cultivation history, plot sizes, and fertilization rates differed between our sites. All these differences can alter PSF effects on microbial community composition and depolymerization capacity. In our case, PSF effects were not found across both fields for every parameter tested. This may be due to differences in pedoclimatic conditions, which alter soil physicochemical properties across both fields (A’Bear et al., 2014). Limitations created by different agricultural practices in our experiment also highlighted the need to investigate PSF effects more deeply. As mentioned above, different faba bean incorporation at Cloutier and Palmarolle certainly influenced microbial communities and their functions. But, even with different residue management, we found that the previous crops’ PSF influenced microbial community composition and soil ammonium content. Alternatively, any of the above-mentioned differences between the two fields could have played a role. It seems, therefore, possible to enhance PSF when combining crop rotations with other agricultural practices.

## 5. Conclusion

In conclusion, even with limitations encountered by doing an experiment under actual farm operating conditions (field size, block number, residue management), we showed that faba bean increased soil ammonium and wheat grain protein content as compared to peas, at Cloutier. We had to reject our hypothesis that this effect on wheat grain protein content was due to changes in potential soil enzymatic activities, as we found little statistical support for it. We have, however, come up with two alternative explanations. First, most significant shifts in microbial communities due to previous legume identity were in fungal ASVs. This could result in a shift in the relative dominance of bacteria and fungi, which, because of differences in C:N ratio between the two groups, could result in differences in mineralization rates. Second, as soil enzymatic activities are substrate-limited, the larger root system of faba bean could have brought more substrate to these plots, increasing mineralization. Depolymerization of cellulose and proteins are widely distributed functions among soil microbial communities, and their *in-situ* activities are probably restricted by substrate availability, whereas the ratio between immobilization/mineralization probably varies with the soil microbial community composition. More field experiments with *in-situ* measurements would be needed to fully understand the role of plant-soil feedback in crop rotation systems to improve our practices and sustainably meet crops’ nutritional needs.

## 7. Acknowledgements

We extend our gratitude to our interns: Raphaël Boisvert, Sopheaktra Honn Kuhn, Marc-Antoine Duchesne, Hubert Côté and Lisa Colleoni whose hard work during field sampling greatly contributed to this project. We are thankful to Johanne Massy, Martin Jacques, Werbson Lima Barroso, Mahefa Ravoavison, Sendy Augustin Salomon and Thierry Nousseassi for their technical assistance, especially with harvest and assessment of wheat yield and protein content. We are also thankful to the farmers who generously provided access to their fields and supported this project. Their collaboration and commitment were essential to the success of this study.

