## Supplemental material for "Previous legume identity influence wheat rhizosphere microbial communities and grain protein content"

Supplementary results

Table S1: PERMANOVA results of Hellinger-transformed Euclidian distances from bacterial communities (16S) data at Cloutier and Palmarolle

|  | Variable | Df | SumOfSqs | R2 | F | p-value | |
| --- | --- | --- | --- | --- | --- | --- | --- |
| Clou  2024  2 sampling | **Fertilization** | **1** | **0.395** | **0.029** | **1.391** | **0.002** | ****** |
|  | **Previous crop** | **1** | **0.378** | **0.028** | **1.333** | **0.006** | ****** |
|  | Compartment | 1 | 0.271 | 0.020 | 0.955 | 0.481 |  |
|  | **Fertilization: Previous crop** | **1** | **0.332** | **0.025** | **1.170** | **0.048** | ***** |
|  | Fertilization: Compartment | 1 | 0.251 | 0.019 | 0.884 | 0.818 |  |
|  | Previous crop: Compartment | 1 | 0.267 | 0.020 | 0.941 | 0.55 |  |
|  | Fertilization: Previous crop: Compartment | 1 | 0.245 | 0.018 | 0.865 | 0.886 |  |
|  | Residual | 40 | 11.353 | 0.841 |  |  |  |
|  | Total | 47 | 13.493 | 1.000 |  |  |  |
| Clou 2024  1 sampling | Fertilization | 1 | 0.248 | 0.041 | 0.934 | 0.540 |  |
|  | Previous crop | 1 | 0.284 | 0.046 | 1.068 | 0.271 |  |
|  | Fertilization : Previous crop | 1 | 0.272 | 0.044 | 1.022 | 0.355 |  |
|  | Residual | 20 | 5.321 | 0.869 |  |  |  |
|  | Total | 23 | 6.125 | 1 |  |  |  |
| Clou 2023  2 sampling | **Previous crop** | **1** | **0.644** | **0.042** | **2.112** | **0.001** | ****** |
|  | **Compartment** | **1** | **0.657** | **0.043** | **2.156** | **0.001** | ****** |
|  | **Previous crop: Compartment** | **1** | **0.581** | **0.038** | **1.908** | **0.001** | ****** |
|  | Residual | 44 | 13.412 | 0.877 |  |  |  |
|  | Total | 47 | 15.294 | 1 |  |  |  |
| Pal 2024  2 sampling | Fertilization | 1 | 0.384 | 0.024 | 1.148 | 0.083 |  |
|  | **Previous crop** | **1** | **0.477** | **0.030** | **1.427** | **0.006** | ****** |
|  | **Compartment** | **1** | **0.454** | **0.029** | **1.360** | **0.017** | ***** |
|  | Fertilization: Previous crop | 1 | 0.381 | 0.024 | 1.140 | 0.104 |  |
|  | Fertilization: Compartment | 1 | 0.296 | 0.019 | 0.885 | 0.714 |  |
|  | Previous crop: Compartment | 1 | 0.304 | 0.019 | 0.909 | 0.633 |  |
|  | Fertilization: Previous crop: Compartment | 1 | 0.263 | 0.016 | 0.786 | 0.984 |  |
|  | Residual | 40 | 13.372 | 0.839 |  |  |  |
|  | Total | 47 | 15.931 | 1.000 |  |  |  |
| Pal  2024  1 sampling | Fertilization | 1 | 0.283 | 0.04 | 0.926 | 0.437 |  |
|  | Previous crop | 1 | 0.289 | 0.042 | 0.945 | 0.379 |  |
|  | Fertilization : Previous crop | 1 | 0.276 | 0.040 | 0.902 | 0.467 |  |
|  | Residual | 20 | 6.112 | 0.878 |  |  |  |
|  | Total | 23 | 6.960 | 1 |  |  |  |
| Pal  2023  2 sampling | **Previous crop** | **1** | **0.475** | **0.029** | **1.358** | **0.041** | ***** |
|  | **Compartment** | **1** | **0.695** | **0.042** | **1.985** | **0.001** | ****** |
|  | Previous crop: Compartment | 1 | 0.369 | 0.022 | 1.053 | 0.291 |  |
|  | Residual | 43 | 15.051 | 0.907 |  |  |  |
|  | Total | 46 | 16.589 | 1 |  |  |  |

*: p<0.05; **p<0.01; ***:p<0.001

Df, SumOfSqs, R2, F and p-value in bold are the significant and/or almost significant values

Table S2: PERMANOVA results on Hellinger-transformed Euclidean distances from fungal communities (ITS) data at Cloutier and Palmarolle

|  | Variable | Df | SumOfSqs | R2 | F | p-value | |
| --- | --- | --- | --- | --- | --- | --- | --- |
| Clou 2024  2 sampling | **Fertilization** | **1** | **0.471** | **0.028** | **1.350** | **0.007** | ******* |
|  | **Previous crop** | **1** | **0.720** | **0.043** | **2.065** | **0.001** | ****** |
|  | **Compartment** | **1** | **0.417** | **0.025** | **1.198** | **0.037** | ***** |
|  | Fertilization: Previous crop | 1 | 0.382 | 0.023 | 1.095 | 0.158 |  |
|  | Fertilization: Compartment | 1 | 0.287 | 0.017 | 0.823 | 0.917 |  |
|  | Previous crop: Compartment | 1 | 0.285 | 0.017 | 0.818 | 0.934 |  |
|  | Fertilization: Previous crop: Compartment | 1 | 0.316 | 0.019 | 0.908 | 0.679 |  |
|  | Residual | 40 | 13.940 | 0.829 |  |  |  |
|  | Total | 47 | 16.817 | 1.000 |  |  |  |
| Clou 2024  1 sampling | Fertilization | 1 | 0.332 | 0.042 | 0.980 | 0.390 |  |
|  | Previous crop | 1 | 0.333 | 0.0427 | 0.985 | 0.392 |  |
|  | Fertilization: Previous crop | 1 | 0.379 | 0.048 | 1.118 | 0.158 |  |
|  | Residual | 20 | 6.774 | 0.866 |  |  |  |
|  | Total | 23 | 7.818 | 1 |  |  |  |
| Pal  2024  2 sampling | Fertilization | 1 | 0.394 | 0.023 | 1.116 | 0.108 |  |
|  | **Previous crop** | **1** | **0.844** | **0.049** | **2.391** | **0.001** | ******* |
|  | **Compartment** | **1** | **0.503** | **0.029** | **1.426** | **0.007** | ****** |
|  | **Fertilization: Previous crop** | **1** | **0.441** | **0.025** | **1.249** | **0.043** | ***** |
|  | Fertilization: Compartment | 1 | 0.296 | 0.017 | 0.839 | 0.824 |  |
|  | Previous crop: Compartment | 1 | 0.396 | 0.023 | 1.123 | 0.123 |  |
|  | Fertilization: Previous crop: Compartment | 1 | 0.314 | 0.018 | 0.891 | 0.673 |  |
|  | Residual | 40 | 14.114 | 0.816 |  |  |  |
|  | Total | 47 | 17.302 | 1.000 |  |  |  |
| Pal  2024  2 sampling | Fertilization | 1 | 0.378 | 0.043 | 1.017 | 0.322 |  |
|  | **Previous crop** | **1** | **0.539** | **0.062** | **1.448** | **0.007** | ****** |
|  | Fertilization:Previous crop | 1 | 0.387 | 0.044 | 1.039 | 0.27 |  |
|  | Residual | 20 | 7.444 | 0.851 |  |  |  |
|  | Total | 23 | 8.748 | 1 |  |  |  |

*: p<0.05; **p<0.01; ***:p<0.001

Df, SumOfSqs, R2, F and p-value in bold are the significant and/or almost significant values
